# H3K36M provokes cellular plasticity to drive aberrant glandular formation and squamous carcinogenesis

**DOI:** 10.1101/2022.08.19.504575

**Authors:** Eun Kyung Ko, Amy Anderson, Jonathan Zou, Sijia Huang, Sohyun Cho, Faizan Alawi, Stephen Prouty, Vivian Lee, Kai Ge, John T. Seykora, Brian C. Capell

## Abstract

Epigenetic dysregulation is pervasive in cancer, frequently impairing normal tissue development and differentiation^1^. Beyond alterations in histone modifying enzymes, “oncohistone” mutations have been described across a variety of cancers^2-4^, although the *in vivo* effects and underlying mechanisms behind these observations have not been well-studied and remain unclear. Here, by inducing the *in vivo* expression of histone H3.3 carrying a lysine to methionine (K to M) mutation at position 36 (H3K36M) in self-renewing stratifying epithelial tissues, we show that the H3K36M oncohistone dramatically disrupts normal epithelial differentiation, leading to extensive tissue dysplasia characterized by a significant increase in mitotic, proliferative basal stem cells. Furthermore, this differentiation blockade promotes increased cellular plasticity and enrichment of alternate cell fates, and in particular the aberrant generation of excessive glandular tissue including both hypertrophic salivary, sebaceous, and meibomian glands. Upon carcinogen stress, H3K36M mice display markedly enhanced squamous tumorigenesis. These aberrant phenotypic and gene expression manifestations are associated with global loss of H3K36me2 and concomitant gain of H3K27me3. Collectively, these results have revealed a previously unknown critical role for H3K36 methylation in both the *in vivo* maintenance of proper epithelial cell fate decisions and the prevention of squamous carcinogenesis. Additionally, they suggest that H3K36 methylation modulation may offer new avenues for the regulation of numerous common disorders driven by over- or under-active glandular function.

---

Histone lysine methylation across the epigenome provides a highly dynamic system to offer precise coordination of virtually all DNA-templated processes^5^. For example, chromatin state can provide the blueprint for the proper establishment and maintenance of developmental trajectories. When dysregulated, alterations in cell fate as well as uncontrolled cancerous proliferation may ensue. Consistent with this, mutations in both chromatin modifiers as well as histones themselves are amongst the most common events in both human developmental disorders and cancers^1^.

Methylation of histone H3 lysine 36 (H3K36) has been shown to play critical roles maintaining proper transcriptional fidelity, splicing, and DNA repair^6-9^. Additionally, modifications such as H3K36 trimethylation (H3K36me3) contribute to both proper DNA and RNA methylation^10-13^. In line with these broad ranging functions, mutations in genes encoding both H3K36 modifiers and multiple histone genes have been described in numerous human cancers^2,4,14-16^, and are particularly abundant in human epithelial tissues which are constantly self-renewing and rely on precise control of gene expression programs. This includes head and neck squamous cell carcinoma (HNSCC), one of the most common and deadly cancers worldwide^17^, which harbors common mutations in H3K36 modifiers (i.e. *NSD1, SETD2*) in addition to H3K36M oncohistone mutations^2^. Despite this, the functional importance of H3K36 methylation and H3K36M have not been modeled previously *in vivo* in epithelial tissues.

## H3K36M expression *in vivo* drives extensive tissue dysplasia and aberrant glandular formation

In this study, we created a murine model expressing a dominant negative mutant version of H3.3K36M (LSL-H3K36M)^18^ only in stratifying epithelia via a keratin 14 (Krt14) Cre transgene (Extended data Fig. 1a) that has a human Krt14 promoter directing expression of Cre recombinase. While these Krt14-Cre(+); H3K36M mice (“H3K36M”) were born normally, gross examination revealed significantly reduced overall body size along with patchy pale pink skin ventral skin (Fig. 1a and Extended data Fig. 1b) and significant reduction in body weight for both sexes in comparison to wildtype (WT) controls (Fig. 1b). Skin, tongue, and esophagus were harvested at 3 weeks for examination. Consistent with previous *in vitro* results in other systems and cell types^4,19,20^, the expression of H3K36M *in vivo* in stratifying epithelia such as the skin epidermis led to significant reductions in both H3K36me2 and H3K36me3 (Extended data Fig. 1c), with H3K36me2 being lost to a greater extent than H3K36me3 (Extended data Fig. 1d).

**Figure 1:**
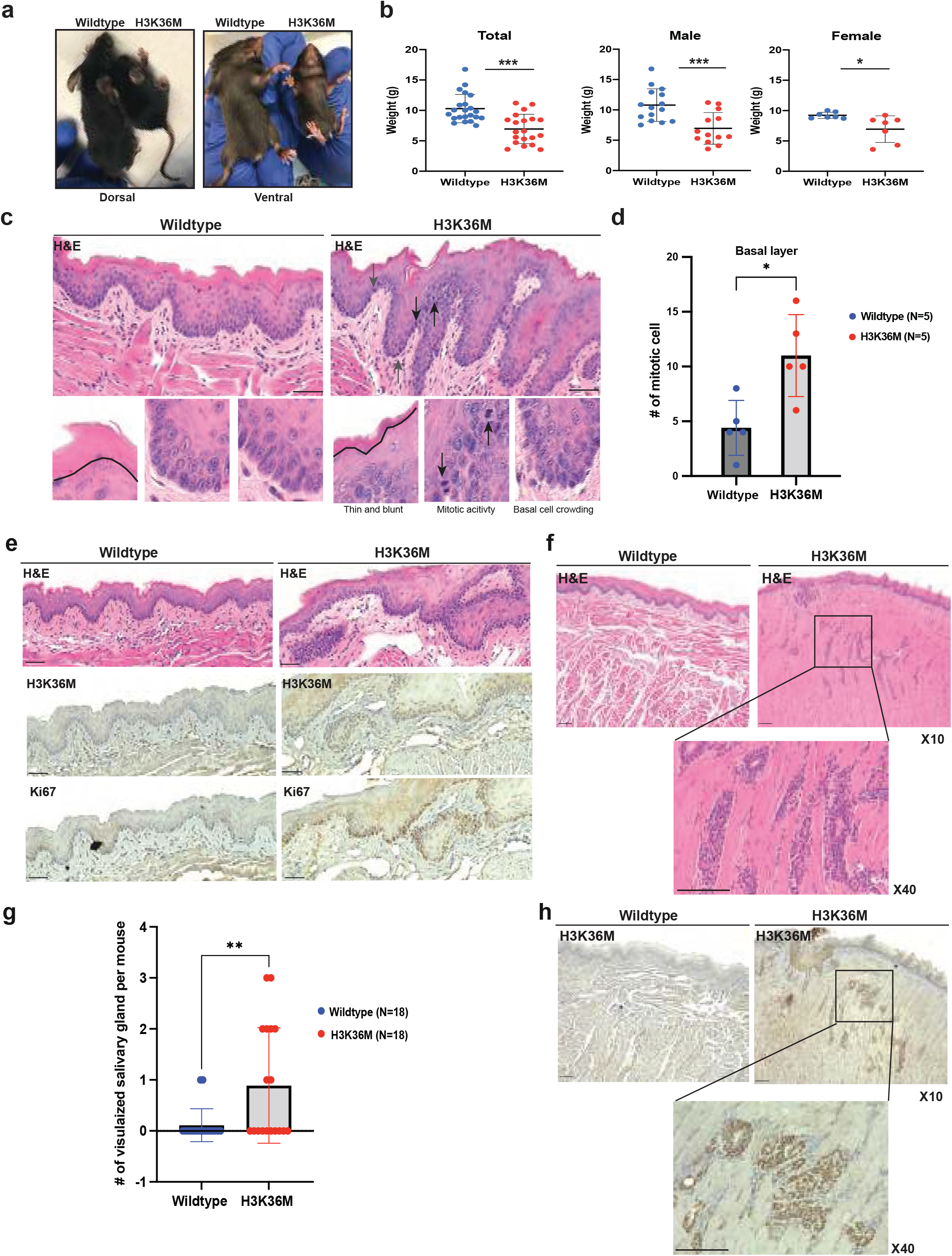
H3K36M expression *in vivo* drives extensive tissue dysplasia and aberrant glandular formation. (a) Representative 3-week-old WT and H3K36M mice. (b) Weights of total (n=20 to 22 per genotype, p<0.001), male (n=13 to 15 per genotype, p<0.001), and female (n=7 per genotype, p<0.05), respectively. (c) H&E staining of 3-week-old WT and H3K36M mice tongues. Black and grey arrows indicate mitotic cells in the basal layer. (d) Counts of the number of mitotic cells in the basal layer from both WT and H3K36M mice (n=5 per genotype, p<0.05). (e) Representative H&E and IHC staining for H3K36M and Ki67 expression. (f) Representative H&E staining of salivary gland in WT versus H3K36M mice. The square inset box indicates that the salivary gland in 10x magnification was further magnified to 40x. (g) Counts of the number of visualized salivary glands per mouse for both WT and H3K36M mice tongues (n=18 per genotype, p<0.01). (h) IHC staining of H3K36M in WT and H3K36M mice tongues demonstrating the strong positivity in the salivary glands. All scale bars are 50um.

Upon Hematoxylin & Eosin (H&E) histology staining, we found that the tongue of H3K36M mice showed features of dysplasia such as increased mitotic activity, basal cell crowding and bulbous drop-shaped rete pegs (Fig. 1c). The number of mitotic cells was significantly greater in H3K36M mice tongue compared to those of the WT mice (Fig. 1d). Specifically, hotspots of mitotic cells were mainly distributed in the dysplastic regions of H3K36M mice tongues. In addition, the expression level of the proliferation marker Ki67 in the basal layer was increased in H3K36M mice tongues. Utilizing an antibody specific for H3K36M suggested that the increased numbers of Ki67 positive cells was enriched particularly in those cells that also stained positive for H3K36M (Fig. 1e). Together, these data suggested that the expression of H3K36M leads to extensive dysplasia and enhanced proliferative activity in the basal cells of the oral epithelium.

Beyond these features of dysplasia, there was evidence of morphological and developmental abnormalities. First, the normal appearance of tongue filiform papillae appeared blunted and reduced in numbers in H3K36M mice (Fig. 1c). Surprisingly, H3K36M mice displayed numerous prominent glandular structures throughout the tongue suggestive of salivary gland hyperplasia that were not observed in control WT mice (Fig. 1f). Whole tongues from mice were harvested under the same conditions regardless genotype and then sectioned longitudinally. This confirmed that few salivary glands were observed in the tongues of WT mice, while numerous aberrant salivary glands were observed in H3K36M mice tongues (Figs. 1f, g).

To test whether this aberrant glandular formation was driven by the expression of the mutant H3K36M protein, we again performed immunohistochemistry (IHC) for the H3K36M protein. While there was no expression of H3K36M in WT mice, H3K36M was strongly expressed in both epithelium and aberrant salivary glands observed in mice carrying the H3K36M allele. Strikingly, the expression of H3K36M was significantly higher in the salivary glands and surrounding connected ducts in comparison even to that of the adjacent oral epithelial tissue in the same H3K36M mouse tongue (Figure 1h).

We next asked whether the expression of the H3K36M protein might lead to aberrant glandular formation in other Krt14-expressing epithelial tissues beyond the tongue. Examination of the skin revealed both larger and greater numbers of sebaceous glands in both the young and aged H3K36M mice dorsal skin (Extended data Fig. 1e). Further detailed examination of tongues from both young and aged H3K36M mice and their controls revealed both types of salivary glands, both serous and mucous, were observed in increasing numbers and with hyperplastic morphology in H3K36M mice tongues (Extended data Fig. 2a). Finally, dissection of the skin of the murine eyelids revealed dramatic increases in the size and number of aberrant meibomian glands (Extended data 2b, c). Collectively, these data strongly support that the expression of H3K36M, an “oncohistone” that disrupts H3K36 methylation, drives global aberrant glandular formation.

## RNA-seq of tongue epithelium reveals a broad loss of differentiation genes and aberrant expression of glandular genes in H3K36M mice

To delineate the transcriptional programs that may be driving the observed phenotypic alterations, we performed RNA-seq on tongue epithelium (dissected from underlying tongue muscle tissue) from WT and H3K36M mice (Fig. 2a). Differential expression analyses revealed that 216 genes were significantly downregulated, while 284 genes were significantly upregulated in H3K36M mice in comparison to WT mice (padj < 0.05, log2FC > +1 or -1)(Figure 2b and Extended Data Table 1). Gene Ontology (GO) analysis demonstrated that significantly downregulated genes suggested an overall loss of expression of genes involved in development, morphogenesis, and differentiation as categories included “Cellular component morphogenesis”, “ERBB4-ERBB4 signaling pathway”, “Cellular developmental process”, “Cell differentiation” and “Epithelium development” were the most enriched categories (Fig. 2b). Amongst the genes making up these categories included *Nrg1*, which stimulates ERBB4 signaling, and *Erbb4* itself, which has been shown to promote oral epithelial differentiation^21,22^. Several downregulated genes also known to be critical for proper oral development included genes involved in Wnt signaling (Fig. 2b, 2c). These included *Bmp4* which has been shown to induce oral ectoderm into dental epithelium and mesenchyme^23^, as well as the Wnt effectors (*Tcf4, Tnik, Fzd3*) and targets (*Flrt3*)^24^. Other major epithelial differentiation associated genes included *Hrnr, Krt10*, and *Ivl*^21^. Notably, HNSCC is more commonly associated with the loss of tumor suppressors than other solid tumors^17^. One such known tumor suppressor is the retinoic acid receptor β (*Rarb*) which we also observed to be significantly downregulated in H3K36M mice. Similarly, a loss of *Tgfbr1* has also been associated with the development and progression of HNSCC^25^. Intriguingly, we also noted the lost expression of two genes intimately involved in tumor immunosurveillance (*Il15, Ctla4*)^26,27^. Collectively, these results demonstrated the mice expressing H3K36M in the tongue epithelium displayed a loss of development and differentiation genes, including the loss of Wnt signaling and numerous genes associated with HNSCC tumor suppression.

**Figure 2:**
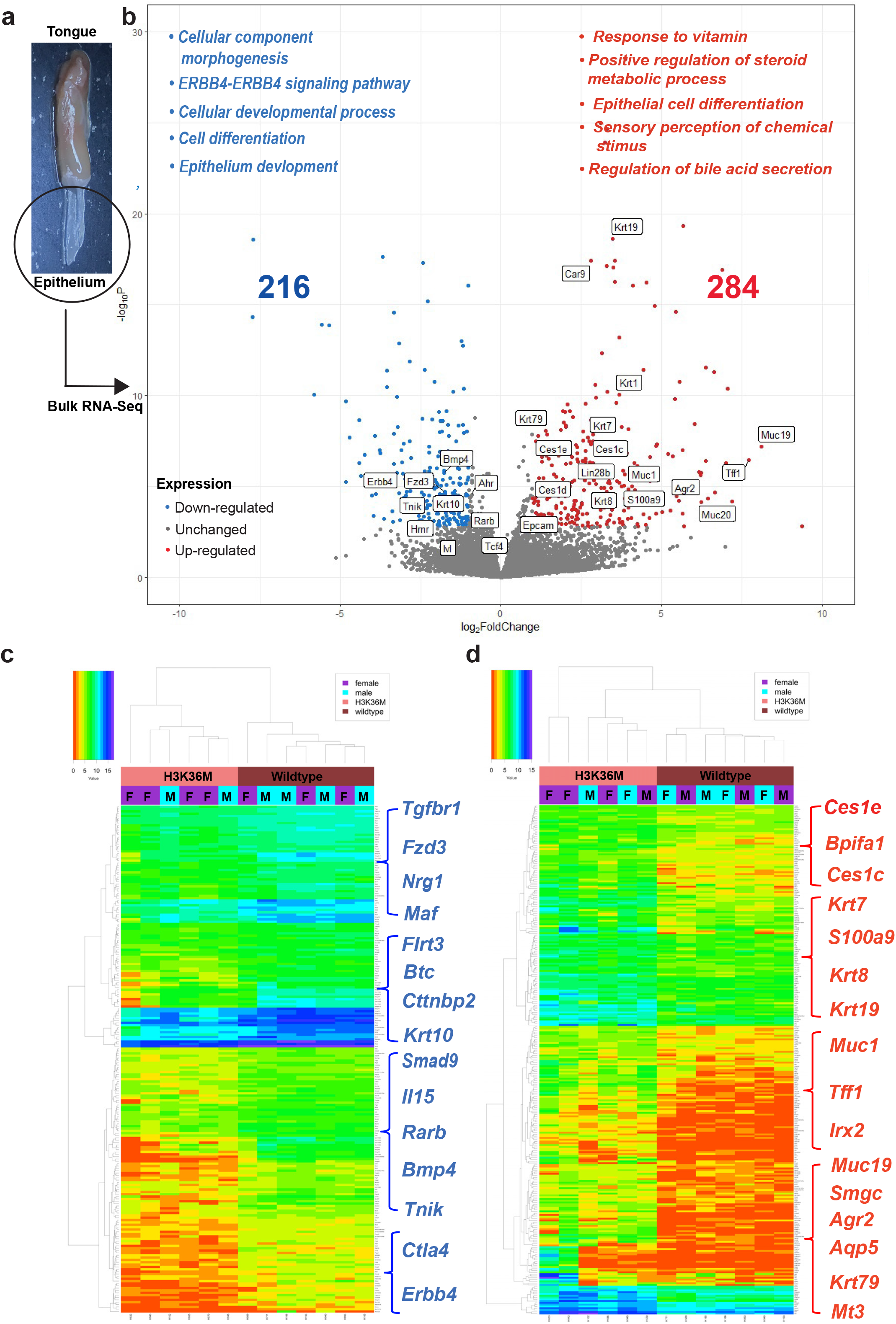
RNA-seq of tongue epithelium reveals a broad loss of differentiation genes and aberrant expression of glandular genes in H3K36M mice. (a) Representative image isolated epithelium from mouse tongue (n=7 WT, n=6 H3K36M, 6 to 8 months of age). (b) Volcano plots representing dysregulated genes in H3K36M mice tongues compared to WT mice tongues along with enriched Gene Ontology (GO) categories for significantly downregulated (blue) and upregulated (red) genes (*Mt3* was removed from heatmap in order to make the rest of the heatmap easier to visualize since it was such an extreme point). (c) Heatmap analysis of genes that were downregulated in H3K36M mice tongues compared to WT mice tongues. (d) Heatmap analysis of genes that were upregulated in H3K36M mice tongues compared to WT mice tongues. M means male and F means female in the heatmap.

Enriched upregulated GO categories included pathways involved secretion and metabolism (Fig. 2b). While “Epithelial cell differentiation” appeared as an upregulated category, it included keratin genes associated with early, primitive epithelial stem cell developmental states (i.e. *Krt7, Krt8, Epcam*) as well as keratin genes that are more specific for glandular epithelium. For example, Krt7 and Krt19 have been shown to be highly enriched in the submandibular glands of the oral epithelium^28^. In a similar fashion, Krt79 has been shown to be specific for sebaceous as well as meibomian, salivary and submandibular glands. Furthermore, blocking Krt79 prevented formation of sebaceous and meibomian glands^29^. Consistent with these findings, several carboxylesterase genes (i.e *Ces1c, Ces1d, Ces1e*)^30^, as well as mucin genes (i.e. *Muc1, Muc19, Muc20*)^31^, known to be expressed in both glandular epithelium and saliva, were found to be enriched in H3K36M mouse tongue. Aquaporin-5 (*Aqp5*), which has been shown to be critical for salivary gland secretory function^32,33^, was also significantly upregulated. *Bpifa1* is a known marker of differentiated salivary glands and was increased over 4-fold in H3K36M mice^31,34^. Numerous other genes known to be abundant and highly expressed in salivary glands that we found upregulated in H3K36M tongue included *Agr2, Tff1, Lpo, Smgc, Krt1, S100a9* and the transcription factors *Ltf* and *Nkx3-1*^31^ (Fig. 2b, 2d).

Importantly, many of these same genes associated with glandular development and activity have also been noted to be increased in HNSCC. For example, high levels of *Krt7* and *Krt19* have been observed in HNSCC^35^, and Krt7 in particular has served as a marker of epithelial mesenchymal transition in some cancers^36^. Likewise, *S100a8* and *S100a9* have also been shown to be upregulated in SCC^37^. Interestingly, metallothionein 3 (*Mt3*) was the most significantly upregulated gene in our dataset and it has been noted to be highly expressed in both SCC^38^, as well as in the salivary gland^39^. Together, these data show that the *in vivo* presence of H3K36M in the oral epithelium causes widespread transcriptional alterations that are marked by a significant loss of differentiation genes along with a striking concomitant upregulation of genes typically expressed in salivary and glandular tissues, as well as genes frequently overexpressed in HNSCC.

## H3K36M mice are more susceptible to squamous carcinogenesis

H3K36M is second most common mutated histone residue overall^3^, and has been observed in a variety of cancers including HNSCC^2,4,40^. Therefore, to better understand the *in vivo* consequences of H3K36M expression in the oral epithelium, we treated mice with 4- Nitroquinoline-1-oxide (4NQO), a water-soluble carcinogen that induces tumors predominantly in the oral cavity. Regular water was given for 20 weeks for the vehicle control groups. For the 4NQO group, 4NQO was given for 12 weeks followed by regular water for an additional 8 weeks (Fig. 3a).

**Figure 3:**
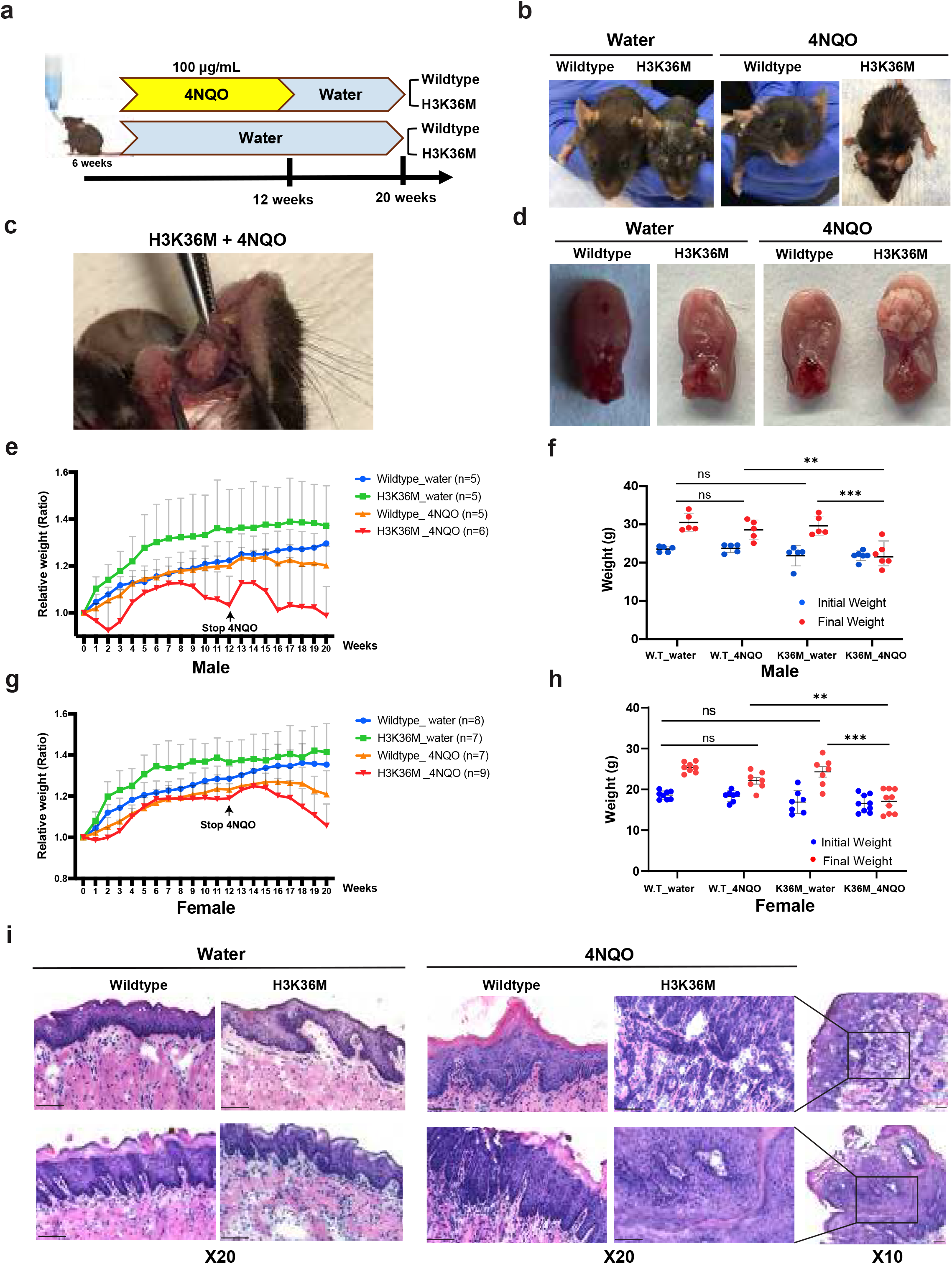
H3K36M mice are more susceptible to squamous carcinogenesis. (a) Schematic diagram of carcinogen exposure study for generating HNSCC using 4NQO. (b) Representative phenotypes per each genotype under either water or 4NQO. (c) Representative image of tumor that generated from H3K36M mouse tongue with 4NQO. (d) Representative images of harvested whole tongue per each genotype under either water or 4NQO. (e) Trends of relative weight that was measured every week for 20 weeks in males. (f) Comparison of absolute weight between initial and final weight in males. (g) Trends of relative weights that was measured every week for 20 weeks in females. (h) Comparison of absolute weights between initial and final weight in females. (i) Representative H&E staining from tongue per each genotype under either water or 4NQO after methanol fixation and cryosectioning. The black square box at 10x was magnified. All scale bars are 50um.

As expected, there were no obvious outward phenotypes observed in WT mice with water. In marked contrast, H3K36M mice receiving water only more aged, with patchy graying, short hair on the skin and a loss of hair around eyelid (Fig. 3b). With 4NQO, all H3K36M mice receiving 4NQO displayed even more severe phenotypes, including smaller body size, more extensive loss of hair on the skin and around the eyelids, as well as less overall movement in comparison to WT mice receiving 4NQO (Fig. 3b). Additionally, visible tumors at the time of dissection at the conclusion of the study were also only observed in H3K36M mice (Fig. 3c, 3d).

During the course of the 20 weeks of the study, all mice were weighed once a week. Relative weight was then calculated by dividing the current weight by the initial weight to observe trends of mouse weight for the water and 4NQO groups. With water, both WT and H3K36M mice became gradually heavier despite the presence of H3K36M. In contrast, with 4NQO, H3K36M mice displayed more variable weight gain during the initial 15 weeks and more significant weight loss during the final 5 weeks of the study in comparison to WT mice receiving 4NQO (Fig. 3e and 3g). The trends of relative weight were similar for both sexes. Interestingly, WT mice did not display any significant difference in weight between the initial weight and final weight with 4NQO, although the WT mice receiving 4NQO groups displayed less weight gain in comparison to WT mice receiving water (Fig. 3f and 3h). In contrast, H3K36M mice demonstrated a significant reduction in weight both in comparison to WT mice both in the vehicle water setting (WT-water vs. H3K36M-water) as well as in the context of receiving 4NQO (i.e. both WT-4NQO vs. H3K36M-4NQO, as well as H3K36M-water vs. H3K36M- 4NQO) (Fig. 3f and 3h). These phenotypic features and weight analyses indicated that H3K36M mice were significantly more impacted by the 4NQO carcinogen in comparison to WT mice.

To better understand the effects of carcinogen exposure, the tongue and esophagus of all mice were harvested for histology, including H&E. As would be expected, no WT mice exposed to water showed any regions of dysplasia, hyperplasia or carcinoma. Both WT mice exposed to 4NQO, and H3K36M receiving water demonstrated regions of squamous hyperplasia and dysplasia, but no evidence of frank carcinoma was observed in any mice (Figure 3i and Extended Data Tables 2 and 3). In striking contrast, all H3K36M mice with 4NQO exhibited various levels of squamous hyperplasia, dysplasia or carcinoma in both sexes (Fig. 3i and Extended Data Tables 2 and 3). Specifically, 44% of H3K36M females (4 out of 9 mice) had either invasive carcinoma or carcinoma in situ while 20% of H3K36M males (1 out 5 mice) had invasive carcinoma. Notably however, we suspect that the lower number of tumors observed in the males was likely secondary to potentially a more severe response to 4NQO administration, as over the course of the study, three H3K36M male mice died sporadically (2 within 2-3 weeks after the provision of 4NQO, and one at 15 weeks after the discontinuation of 4NQO) in comparison to just one female H3K36M mice and no WT mice exposed to 4NQO).

In addition to these effects in the oral epithelia, 4NQO administration has also been known to induce tumor formation in esophagus. While some of esophagi were not able to be observed with H&E staining due to unsuccessful harvesting, the results from 4NQO were the similar to those seen in the tongue (Extended Data Fig. 3). No WT mice receiving water displayed any features of hyperplasia, dysplasia or carcinoma. H3K36M mice receiving water, as well as WT mice exposed to 4NQO showed features of either hyperplasia or dysplasia but no carcinoma was observed regardless of sex. The esophagi of H3K36M mice exposed to 4NQO was once again more severely affected, although the only full esophageal carcinoma was observed in one H3K36M mouse receiving 4NQO (Extended Data Tables 2 and 3). Together, these data strongly support the conclusion that impairment of H3K36 methylation enhances HNSCC squamous tumorigenesis both with and without carcinogen exposure.

## H3K36M drives global concomitant *in vivo* loss of H3K36me2 and gain of H3K27me3 that is spatially associated with aberrant pathological phenotypes and gene expression

Our data suggested that the aberrant expression of H3K36M locked in a more proliferative, stem cell-like state that predisposed the cells to both alternative fates (gland formation) and carcinogenesis (SCC). To better understand the mechanisms behind these phenotypic manifestations, we examined global levels of H3K36 mono-, di-, and trimethylation in the murine tissue in the context of *in vivo* H3K36M expression. Importantly, IHC demonstrated that there was significantly less expression of H3K36me2 in H3K36M mice, and this was specifically apparent in cells expressing higher levels of H3K36M even within the same tissue section (Fig. 4a). H3K36me3 was modestly diminished in H3K36M mice, albeit less so in comparison to H3K36me2 (Fig. 4a). Consistent with previous *in vitro* results, regions of concomitant H3K36M expression and reduced H3K36me2 displayed strong increases in the repressive modification, H3K27me3 (Fig. 4a), which is known to accumulate aberrantly in regions of H3K36me2 loss^19,20,41^.

**Figure 4:**
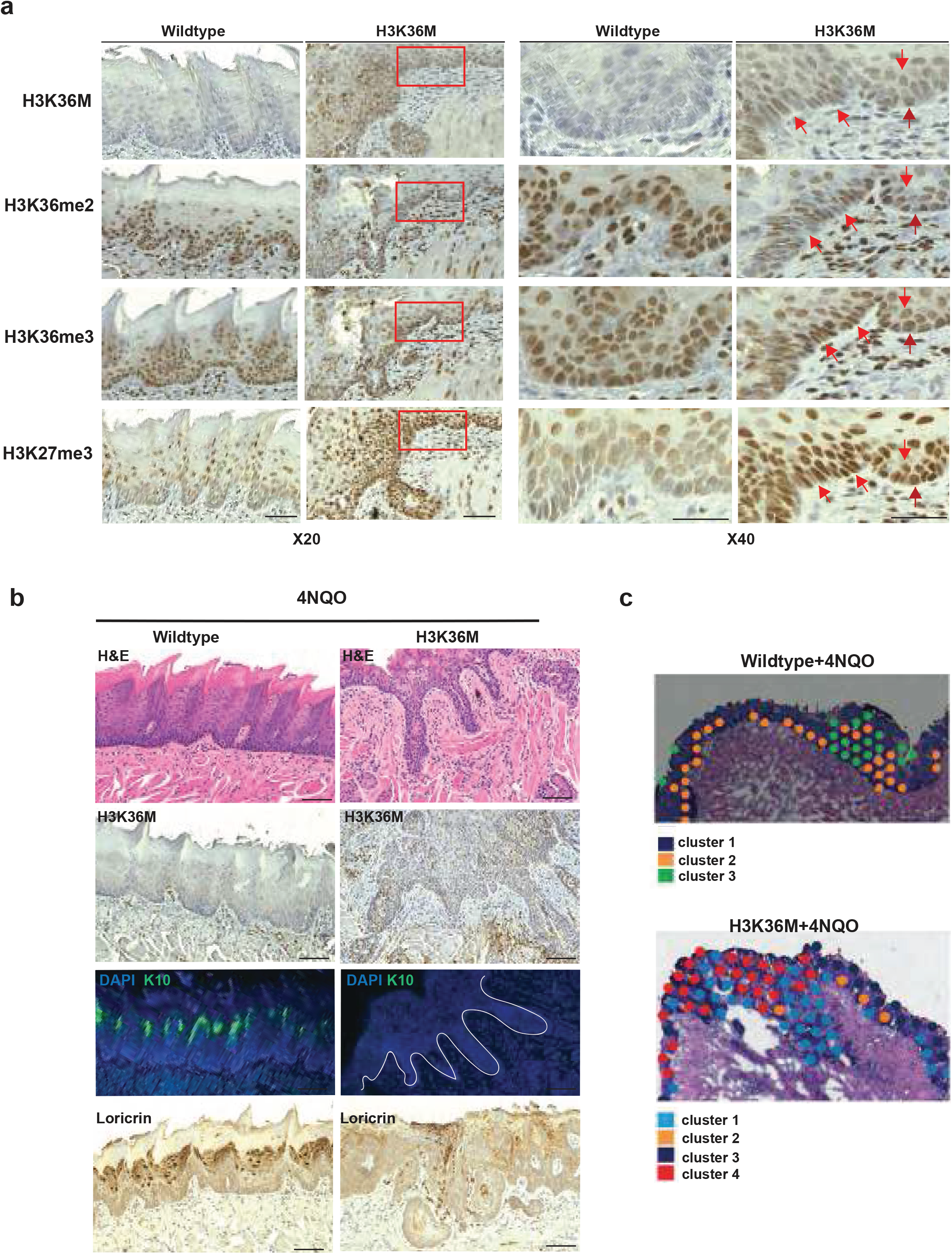
H3K36M drives global concomitant *in vivo* loss of H3K36me2 and gain of H3K27me3 that is spatially associated with aberrant pathological phenotypes and gene expression. IHC staining for the expression levels of H3K36M, H3K36me2, H3K36me3 and H3K27me3 from both WT and H3K36M mice tongues. Red square boxes at 20x were further magnified to 40x. Red arrows indicate differing expression levels of the aforementioned target proteins of interest in the same cell. (b) IHC and IF staining for expression levels of H3K36M, keratin 10 (K10), and Loricrin from both WT and H3K36M mice tongues with 4NQO for 20 weeks. (c) Representative images from Loupe browser following spatial transcriptomic analysis from WT and H3K36M exposed to 4NQO (spot size 55um).

Given the ability of H3K27me3 to directly suppress epithelial differentiation genes^42^, these results supported a model whereby the accumulation of H3K36M led to H3K27me3 accumulation and the suppression of epithelial differentiation *in vivo*. In further support of this, direct staining of H3K36M tongue tissue revealed a complete loss of epithelial differentiation-associated proteins such as keratin 10 (Krt10) and loricrin (Fig. 4b). To more directly explore how the transcriptional alterations driven by H3K36M accumulation contributed to phenotypic manifestations, we performed spatial transcriptomics on WT and H3K36M mice upon 4NQO carcinogen exposure. Here, our results suggested that, while both WT and H3K36M mice possessed more and less differentiated epithelial regions within hyperplastic/dysplastic tissue, only H3K36M mice revealed a unique cluster of cells which displayed both a loss of oral epithelial differentiation genes (i.e. *Krt13, Lor, Lce3a, Lce3b, Lce3c, Lce3f, Cdsn*), along with an aberrant upregulation of genes associated with ductal and glandular epithelia (i.e. *Krt7, Krt19, Ltf*) (Fig. 4c and Extended Data Tables 4 and 5). Interestingly, human HNSCC’s carrying mutant H3K36M have also been shown to express reduced levels of differentiation genes, such as similar late cornified envelope (i.e. *LCE3B, LCE3D, LCE3E*) genes observed in our murine model^2^. Together, these observations suggest that H3K36M leads to significant disruption of the chromatin landscape, and in particular a concomitant loss of H3K36me2 and gain of repressive H3K27me3 that likely suppresses normal epithelial differentiation, in turn creating epigenetic plasticity that provokes aberrant gland formation and a more permissive environment for carcinogenesis.

## Discussion

The disruption of H3K36 methylation in the form of oncohistone mutations like H3K36M has been described in human cancers (including HNSCC) and modeled *in vitro* in various cellular systems^2,4,19,20^. In addition, one previous study found that the global induction of H3K36M in mice led to defects in hematopoiesis, severe anemia, and premature death^43^, while another study utilizing tissue-specific Cre drivers demonstrated that disruption of H3K36 methylation led to differentiation defects during adipogenesis and myogenesis^18^. Here we provide the first *in vivo* model of H3K36M in stratifying epithelial tissues. Collectively, our data demonstrate not only how these mutations create a more permissive, less differentiated state that predisposes to squamous carcinogenesis, but unexpectedly, also uncover a previously unknown role for H3K36 methylation in directing proper epithelial cell trajectories to prevent aberrant glandular formation.

Our results show directly that individual cells expressing H3K36M accumulate aberrant H3K27me3 at the expense of H3K36me2, and to a lesser extent, H3K36me3, and that these cells are particularly enriched in regions of pathological aberrant gland formation and/or squamous hyperplasia, dysplasia, and carcinoma. Consistent with the observed pathological glandular changes seen in H3K36M mice, we observe a striking enrichment of a salivary gland gene expression in the setting of lost oral epithelial differentiation genes. Beyond the ability of H3K36M to disrupt the balance of H3K36me2/3 and H3K27me3 as observed in our study, given the ability of H3K36 methylation to affect both RNA and DNA methylation, we anticipate that future studies can further investigate how the presence of H3K36M in epithelial tissues may disrupt the recruitment of other epigenetic modifiers known to interface with H3K36, including DNMT3A, DNMT3B, and METTL14^10-13^.

Indeed, while the formation of these hyperplastic and excessive salivary, meibomian, and sebaceous glands can be considered pathological in these mice, it suggests a possibility whereby epigenetic manipulation may be able to serve as a therapeutic approach towards either provoking or suppressing gland formation in pathological conditions ranging from Sjogren’s syndrome and dry eye disease to acne and sebaceous hyperplasia.

## EXTENDED DATA

**Extended data figure 1:**
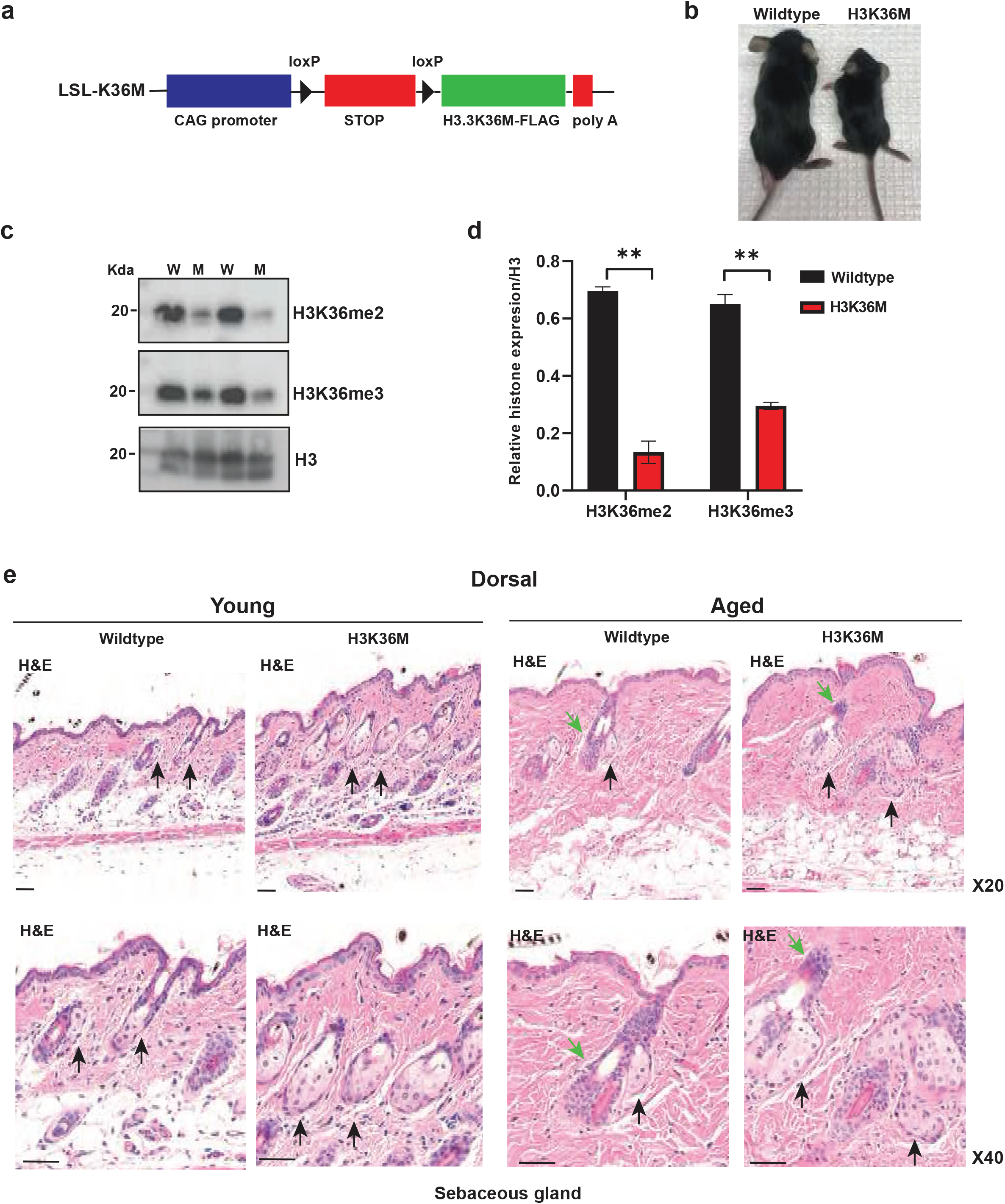
(a) Schematic structure of the LSL-K36M transgenic construct. (b) Representative phenotype of 3-week-old mice from WT and H3K36M genotypes. (c) Immunoblots of H3K36me2, H3K36me3 and total histone H3 isolated from skin epidermis from WT and H3K36M mice. (d) Quantification of relative histone modification expression for H3K36me2 and H3K36me3. (e) H&E staining of sebaceous glands for both young and aged dorsal skin from each genotype. Black arrow indicates the sebaceous gland and green arrow indicates the hair follicle. All scale bars are 50um.

**Extended data figure 2:**
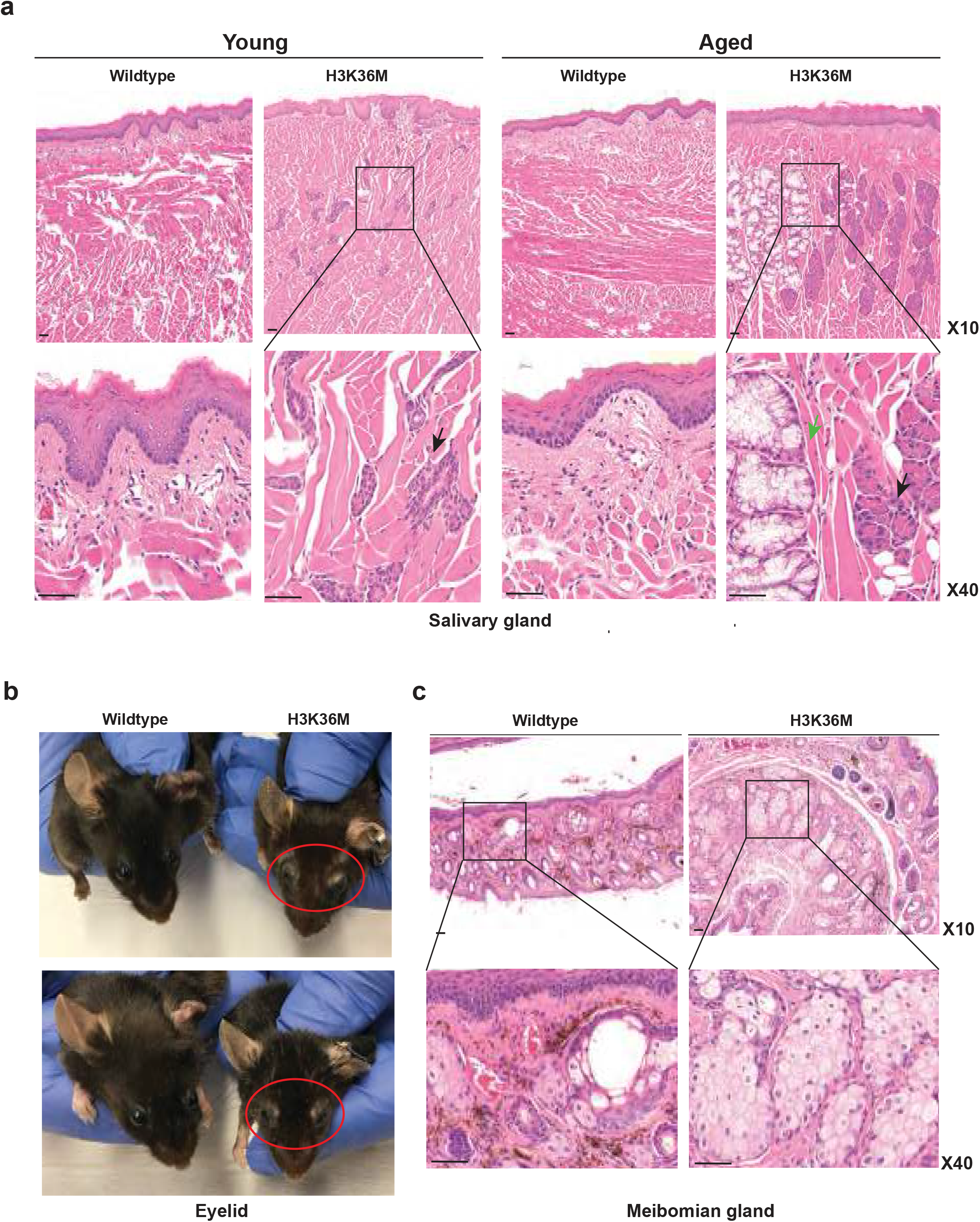
(a) H&E images of salivary glands for both young and aged mice from each genotype (WT vs. H3K36M). Salivary gland in square box at 10x was magnified. Green arrow indicates the mucous salivary gland and black arrow indicates the serous salivary gland. Representative images of the eyelid from both aged WT and H3K36M mice. (c) H&E images of meibomian glands for both aged WT and H3K36M mice. Square box that indicates the meibomian gland at 10x magnification was further magnified to 40x. All scale bars are 50um.

**Extended data figure 3.**
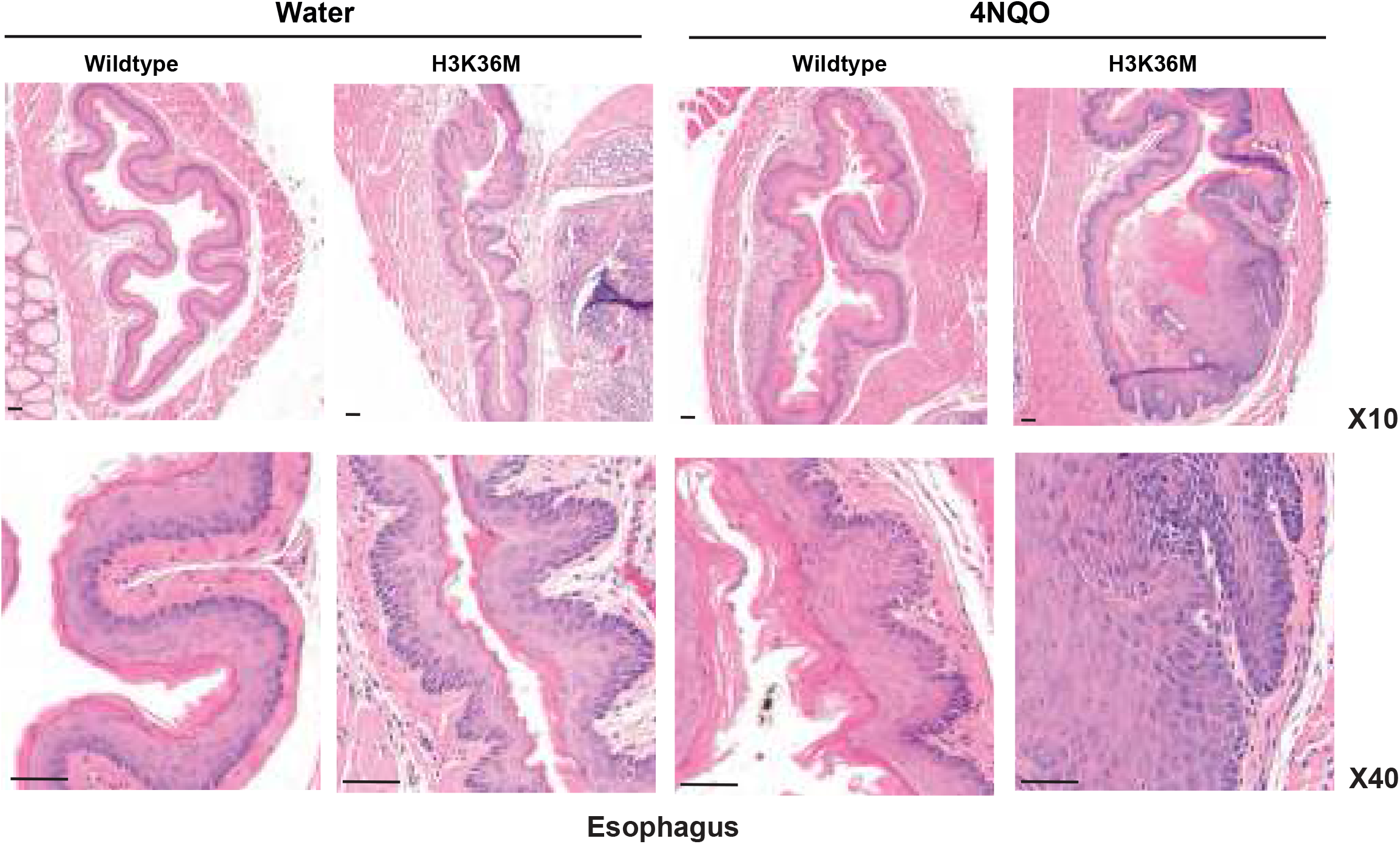
Representative H&E staining from esophagus per each genotype under either water or 4NQO after 4% PFA fixation. Black arrows at 10x were magnified to 40x. All scale bars are 50um.

**Extended data table 1. DESeq2 differential expression analysis of WT versus H3K36M mice tongue epithelia**.

**Extended data table 2. Data on 4NQO induced oral and esophageal cancer development in female mice**. The number in parenthesis was the number that was examined. H is for hyperplasia, D is for dysplasia, and T is for tumor.

**Extended data table 3. Data on 4NQO induced oral and esophageal cancer development in male mice**. The number in parenthesis was the number that was examined. H is for hyperplasia, D is for dysplasia, and T is for tumor.

**Extended data table 4. Spatial transcriptomics run metrics**.

**Extended data table 5. Spatial transcriptomics WT-4NQO cluster analysis**.

**Extended data table 6. Spatial transcriptomics WT-4NQO cluster analysis**.

## METHODS

### Generation of H3K36M mice

LSL-K36M (H3K36M) transgenic mice were previously described^18^. The mice carried full-length H3.3 with a K36M point mutation and a C-terminal FLAG tag fused downstream of the CAG promoter with loxP-STOP-loxP cassette in the middle (Extended data Fig. 1a). H3K36M mice were crossed with Krt14-Cre mice (Jackson Labs #005107) to generate Krt14-Cre; H3K36M mice. Polymerase chain reaction (PCR) was done using a Thermo Phire Animal Tissue PCR Kit (F140WH) for genotyping using the following primers: H3K36M (forward: CTAGCTGCAGCTCGAGTGAACCATGGC and reverse: TTCGCGGCCGCGAATTCCTAGGCGTAGTCG), and Cre (forward: GAACCTGATGGACATGG and reverse: AGTGCGTTCGAACGCTAGAGCCTGT). All mouse experiments were performed in accordance with the University of Pennsylvania Guide for the Care and Use of Laboratory Animals and approved by the Institutional Animal Care and Use Committee of the University of Pennsylvania. Mice were maintained on a 12 h light/dark cycle and stand pellet with regular water, unless otherwise indicated during treatment with the water-soluble carcinogen, 4-Nitroquinoline-1-oxide (4NQO).

### Weight collection and statistical analyses

Following sacrifice, mice at 3-4 weeks of age were measured for total body weight. An unpaired t-test was used to calculate significance between two groups (WT vs. H3K36M) when considered for genotype only (Fig. 1b). During the 4NQO study, mice body weight was assessed weekly without euthanasia. A one-way analysis of variance (ANOVA), followed by Tukey’s multiple comparison test was used to calculate significance difference between 4 groups (Figs. 2f, 2h). All data were analyzed using GraphPad Prism version 5.03 (GraphPad Software, Inc. CA, 92037 USA). A value of *p* < 0.05 was considered statistically significant. Data are presented as mean ± standard error of the mean (SEM).

### Histology

Mouse dorsal skin, ventral skin, tongue, esophagus, and eyelid tissue were harvested for histological examination by Core A of the Penn Skin Biology and Disease Resource-based Center (SBDRC). Hematoxylin & Eosin (H&E) staining was processed by the Penn SBDRC Core A. A Keyence (Keyence, BZ-X710) microscope was used to capture representative images. Exposure times (10x for 250 seconds, 20x for 80 seconds, 40x for 25 seconds) and intensity were kept constant for all mouse samples.

### RNA extraction from mouse tongue epithelium

Isolation of the epithelium from mice (Fig. 2a) was conducted with instruction from previous paper with minor modifications^44^. Briefly, 5 units of dispase (Corning®, #354235) and 10mg/ml of collagenase (Gibco, #17018-029) in 1X PBS were injected into the area beneath the tongue epithelium using a 1ml syringe (BD, #309626) utilizing a microscope (Olympus SZ61) for visualization. The injected tongue was then wrapped using plastic wrap after injection to prevent evaporation and incubated for 22 mins at 37□. The epithelium was separated using sharp forceps under the microscope and then suspended in Trizol LS (Ambion, #10296028) for RNA extraction on ice. The epithelium in Trizol was then transferred into Lysing Matrix D (MP Biomedicals, #6913100) and homogenized using program 1 of FastPrep-24 homogenizer (MP Biomedicals). The supernatant after centrifugation was transferred to a new tube and RNA was extracted using RNeasy Mini Kit (Qiagen, #74106). All procedures for RNA extraction were performed following the manufacturer’s instructions.

### RNA-sequencing

RNA-sequencing (RNA-seq) was performed as described previously^45^. Briefly, all RNA-seq libraries were prepared using the NEBNext poly(A) mRNA magnetic isolation module (NEB, #E7490L) followed by NEBNext Ultra Directional RNA library preparation kit (NEB, #E7920L) for Illumina. Library quality was checked by an Agilent BioAnalyzer 2100 and libraries were quantified using the Library Quant Kit® (NEB, #E7630L) for Illumina. Libraries from WT or H3K36M mice were sequenced on the NextSeq500 platform (75-base-pair (bp) single-end reads). Read alignment was performed using Illumina RNA-seq alignment workflow (version 2.0.1). Briefly, reads were mapped to *Mus musculus* University of California at Santa Cruz (UCSC) mouse GRCm38/mm10 reference genome using RefSeq annotation with STAR aligner (version 2.6.1a)^46^. Differential expression analysis was performed with DESeq2 on *Mus musculus* UCSC GRCm38/mm10 reference genome using Illumina RNA-seq differential expression analysis workflow (version 1.0.1)^47^. Among 27914 total genes, 500 were successfully tested for differential expression analysis. At Padj-value <=0.05 (log2FC >+1.0 or -1.0), 216 genes were downregulated whereas 284 genes were upregulated in the H3K36M mice tongues compared to WT tongues (n=7 WT, and n=6 H3K36M mice).

### RNA-seq heatmap analysis

We generated the differential gene expression correlation heatmap with hierarchical clustering, where Euclidean distance measure was employed. From the DESeq2 differential expression analysis results, we further filtered the genes with adjusted padj-values smaller than 0.05 and absolute log fold change greater than 1. Gene expression counts data are log2 (normalized counts +1) transformed and depicted from red to blue (low to high expression) in the heatmap.

### 4-Nitroquinoline 1-oxide (4NQO) study

A 4NQO carcinogen exposure study was performed as described previously^48^, with slight modifications for our mouse model. In the 4-Nitroquinoline 1-oxide (4NQO) study, 6 weeks old mice in the 4NQO group were given 100ug/ml of 4NQO (Sigma-Aldrich, #N8141), a water soluble chemical carcinogen, in propylene glycol (Sigma-Aldrich, #P4347) for 12 weeks. After stopping 4NQO, the 4NQO groups were transitioned to regular water for 8 weeks. In the vehicle control groups, 6 weeks old mice were given with regular water for 20 weeks. The concentration and duration of the 4NQO administered were chosen based on pilot data in our lab utilizing the same mouse strain. Four groups (WT-water, WT- 4NQO, H3K36M-water, and H3K36M-4NQO) were utilized in both male and female mice (Fig. 3a). All mice were weighed once a week during the 20 week experiment. The tongue was embedded using an Optimal Cutting Temperature compound (OCT) (Tissue-Tek, #4583) for frozen sectioning compatible with spatial transcriptomics analysis.

### Spatial transcriptomics library preparation and sequencing

For spatial transcriptomics, Spatial Gene Expression slides (10x Genomics, #1000188) were used, following the manufacturer’s instructions. Briefly, freezing and embedding of the tongues from the 4NQO were performed separately. Tongues were embedded in cryomold (Fisher, #22363552) using chilled OCT compound on powdered dry ice following freezing in 2- Methylbutane (Sigma, #270342) and a dry ice bath. The frozen embedded tissues were cryosectioned in a cryostat (MICROM, HM505E) with 10um thickness. The frozen slides were fixed in pre-chilled methanol at -20□for 30 min and H&E staining was performed following manufacturer’s instructions. All Capture Areas in the slide following H&E staining were captured individually using the Keyence (Keyence, #BZ-X710) with the bright field images setting at 20x magnification for visualization. The fixed and stained gene expression slide was treated with a permeabilization enzyme specific for mouse tongue determined by the Spatial tissue optimization slide kit (10x Genomics, #1000191) for 24 minutes on the thermal cycler (Bio-Rad, 1851197). Synthesis of cDNA and second strand were performed using reverse transcriptase and second strand reaction (primer and enzyme). Real-time PCR (Applied Biosystems, #ViiA 7) was performed using the KAPA SYBR FAST qPCR master Mix (KAPA Biosystems, #KK4600) to obtain the proper number of PCR cycle needed to amplify of cDNA. Finally, 16 cycles were used for cDNA amplification and 1ul of amplified cDNA was analyzed with High Sensitivity DNA Chip (Agilent Technologies, #5067-4626) using a Bioanalyzer (Agilent Technologies, #2100) to calculate the proper number of PCR cycles in library construction preparation. Twenty five percent (10u1) of amplified cDNA was used for library construction. Dual index kit TT Set A (PN-1000215) was used for a multiplexed sequencing.

PCR with 19 cycles were performed for the libraries using the dual index primers. High Sensitivity DNA Chip (Agilent) and the Library Quant Kit® (NEB, #E7630L) were used for fragmentation and quantification of libraries before sequencing. Libraries were sequenced on the NextSeq500 platform (paired-end reads, 150 cycles) with standalone mode and running parameters with 28 cycles for Read 1, 10 cycles for i7 index, 10 cycles for i5 index, 90 cycles for Read 2.

### Spatial transcriptomics mapping and alignment pipeline

The output data of each sequencing run (Illumina BCL files) were processed through the 10x Genomics Space Ranger (v1.3.1) pipelines. Samples were demultiplexed into FASTQ files using Space Ranger’s mkfastq pipeline. Spatial images were then manually curated with selection of epidermis spots. Space Ranger’s count pipeline was used to align FASTQ files with processed images, detect barcode/unique molecular identifier (UMI) counting, and map reads to the mouse reference transcriptome (Gencode vM23 and GRCm38). The Space Ranger pipeline generates, for each sample, feature-barcode matrices that contain raw counts and places barcoded spots in spatial context on the slide image (cloupe files). Gene expression with spatial information can then be visualized by loading cloupe files onto the Loupe Browser (v6.0.0, 10x Genomics). The Space Ranger output summary metrics can be found in (Extended data Table 4).

### Histone protein extraction from epithelium of mouse skin and immunoblotting

Mouse skin was treated with 1x dispase in PBS at 37□for 45 min. The epidermis was dissociated from the dermis using a fresh scalpels with stainless steel surgical blades (Bard- Paker^®^, #371610) and transferred the epidermis to Lysing Matrix D (MP biomedicals, #6913100). For histone protein extraction, the EpiQuikTM Total Histone Extraction Kit (Epigentek, #OP-0006) was used, following the manufacturer’s instructions. Briefly, 1x pre-lysis buffer was added to the epidermis in the Lysing Matrix D tube and program 1 of FastPrep-24 homogenizer (MP Biomedicals) was used for homogenization of the epidermis. The supernatant was removed after centrifugation and the epidermis was lysed in lysis buffer and incubated on ice for 30 mins. The supernatant fraction was transferred into a new tube after centrifugation and 0.3 volume of balance-DTT buffer was added into the supernatant fraction. For immunoblotting of anti-H3K36me1 (Abcam, #ab9048), anti-H3K36me2 (Abcam, #ab9049) and anti-H3K36me3 (Abcam, #ab9050), 700ng of histone protein and 0.2 um pore PVDF membrane (BioRad, #162- 0186) were used. These three types of H3K36 were normalized by an anti-Histone H3 antibody (Abcam, #ab176842). Blotting images were taken using ChemiDocTM imaging System (Biorad) following ECL prime western blotting detection reagent (Amersham, #RPN2232).

### Immunohistochemistry (IHC) staining

IHC staining was performed as described previously^49^. Briefly, mouse tissue slides were baked for 1 hour at 65°C, deparaffinized in xylene, and rehydrated through a series of graded alcohols. After diH_2_O washes, slides were boiled with antigen unmasking solution (Vector Laboratories, #H-3300) using microwave for antigen retrieval. A VECTASTAIN® Elite® ABC Universal Kit Peroxidase (Vector Laboratories, #PK-6200) was used to detect binding of primary antibodies. All procedures were performed according to the manufacturer’s instruction. Slides were treated with blocking solution in the kit for 1 hour at room temperature (RT) and then were incubated with primary antibodies at 4□overnight. The next day, the slides were treated with biotinylated secondary anti-mouse/rabbit antibody for 90 min and then treated with ABC reagent (Avidin- Biotin Complex) for 30 min at RT following blocking of endogenous peroxidases using 3% hydrogen peroxide in MeOH for 10 min at RT. The staining was visualized with 3,3′- diaminobenzidine (Vector Laboratories, #SK-4100, DAB) as peroxidase substrate. Exposure times for DAB were the exact time within an antibody group. All slides were counterstained with hematoxylin (Vector Laboratories, Hematoxylin QS, #H3404) for 1-2 min at RT, dehydrated in ethanol, cleared in xylene, and mounted with VectaMount (Vector Laboratories, #H-5000).

Primary antibodies, anti-Histone H3, K36M mutant (EMD Millipore, ABE #1447, 1:150), anti- H3K36me2 (Abcam, #ab9049, 1:500), anti-H3K36me3 (Abcam, #ab9050, 1:500), anti- H3K27me3 (Cell Signaling, #9733S, 1:400), anti-Ki-67 (Abcam, #ab16667, 1:500), anti-Loricrin (Abcam, #ab85697,1:500), were used for IHC.

### Immunofluorescence staining

Baking, deparaffinization and rehydration of mouse tissue slides were processed with the same protocol that we used for IHC staining. Slides were treated with 1x Target Unmasking Fluid (TUF)(Pan Path, #Z000R.0000) at 90□for 10 min on the hot plate with stirring for antigen retrieval. The beaker with the slides in TUF was removed from the hot plate and leave to cool to 50LJ. The slides were incubated with BlockAid™ blocking solution (Invitrogen, #B107100) for 2 hours at RT and incubated with primary antibody at 4□overnight. The next day, the slides were washed three times for 10 min with 1x TBST (Cell Signaling, #9997S) with shaking on each wash and were incubated with secondary antibody for 1 hour at RT. The slides were stained for DAPI and mounted with ProLong™ Gold antifade reagent with DAPI (Invitrogen, #P36935) following washing with three times for 10 min with 1x TBST with shaking. Anti-keratin 10 (Abcam, #ab76318, 1:500) for primary antibody and Alexa Fluor 488 donkey anti-rabbit IgG (H+L)(Invitrogen, #A21206) for secondary antibody were used. Pictures were taken with a fluorescence microscope (Leica biosystems, LASX).

### Count of mitotic cells and salivary glands

Mitotic cells in basal layer of mouse tongues were counted as the number of mitoses per 10 high-power fields (HPF)(400x magnification) for either WT or H3K36M mice^50^, respectively. The number of visualized salivary gland was counted from both WT and H3K36M mice. An unpaired t-test was used to assess significance between two groups and a value of *p* <0.05 was considered statistically significant.

### Statistical analyses

All statistical analyses were performed using R or GraphPad Prism 8. Details of each statistical test are included in Materials and Methods. Sample sizes and *P* values are included in the figure legends or main figures.

## ACKNOWLEDGEMENTS

This work was supported by the National Institute of Arthritis and Musculoskeletal and Skin Diseases (NIAMS) (R01 AR077615), the Damon Runyon Cancer Foundation, and the Dermatology Foundation, all to B.C.C. E.K.K. has been supported by a Sun Pharma Society of Investigative Dermatology Innovation Award. This project has also received critical support from the Penn Skin Biology Disease Research Center (P30 5P30AR069589-07).

## AUTHOR CONTRIBUTIONS

E.K.K., A.A., J.Z., S.C. and S.P. performed all experiments. S.H. performed all bioinformatic analysis. S.P., F.A., V.L., and J.T.S. assisted with histological analyses. K.G. provided key reagents. E.K.K. and B.C.C. conceived of this work and wrote the manuscript. All authors have read and approved of the final manuscript.

## REFERENCES

1 Zhao, S., Allis, C. D. & Wang, G. G. The language of chromatin modification in human cancers. Nat Rev Cancer 21, 413–430, doi:10.1038/s41568-021-00357-x (2021).

2 Papillon-Cavanagh, S. et al. Impaired H3K36 methylation defines a subset of head and neck squamous cell carcinomas. Nat Genet 49, 180–185, doi:10.1038/ng.3757 (2017).

3 Nacev, B. A. et al. The expanding landscape of ‘oncohistone’ mutations in human cancers. Nature 567, 473–478, doi:10.1038/s41586-019-1038-1 (2019).

4 Lu, C. et al. Histone H3K36 mutations promote sarcomagenesis through altered histone methylation landscape. Science 352, 844–849, doi:10.1126/science.aac7272 (2016).

5 Husmann, D. & Gozani, O. Histone lysine methyltransferases in biology and disease. Nat Struct Mol Biol 26, 880–889, doi:10.1038/s41594-019-0298-7 (2019).

6 Li, J., Ahn, J. H. & Wang, G. G. Understanding histone H3 lysine 36 methylation and its deregulation in disease. Cell Mol Life Sci 76, 2899–2916, doi:10.1007/s00018-019-03144-y (2019).

7 Rogawski, D. S., Grembecka, J. & Cierpicki, T. H3K36 methyltransferases as cancer drug targets: rationale and perspectives for inhibitor development. Future Med Chem 8, 1589–1607, doi:10.4155/fmc-2016-0071 (2016).

8 Wagner, E. J. & Carpenter, P. B. Understanding the language of Lys36 methylation at histone H3. Nat Rev Mol Cell Biol 13, 115–126, doi:10.1038/nrm3274 (2012).

9 Lam, U. T. F., Tan, B. K. Y., Poh, J. J. X. & Chen, E. S. Structural and functional specificity of H3K36 methylation. Epigenetics Chromatin 15, 17, doi:10.1186/s13072-022-00446-7 (2022).

10 Huang, H. et al. Histone H3 trimethylation at lysine 36 guides m(6)A RNA modification co-transcriptionally. Nature 567, 414–419, doi:10.1038/s41586-019-1016-7 (2019).

11 Weinberg, D. N. et al. The histone mark H3K36me2 recruits DNMT3A and shapes the intergenic DNA methylation landscape. Nature 573, 281–286, doi:10.1038/s41586-019-1534-3 (2019).

12 Dhayalan, A. et al. The Dnmt3a PWWP domain reads histone 3 lysine 36 trimethylation and guides DNA methylation. J Biol Chem 285, 26114–26120, doi:10.1074/jbc.M109.089433 (2010).

13 Baubec, T. et al. Genomic profiling of DNA methyltransferases reveals a role for DNMT3B in genic methylation. Nature 520, 243–247, doi:10.1038/nature14176 (2015).

14 Campbell, J. D. et al. Genomic, Pathway Network, and Immunologic Features Distinguishing Squamous Carcinomas. Cell Rep 23, 194–212 e196, doi:10.1016/j.celrep.2018.03.063 (2018).

15 Cerami, E. et al. The cBio cancer genomics portal: an open platform for exploring multidimensional cancer genomics data. Cancer Discov 2, 401–404, doi:10.1158/2159-8290.CD-12-0095 (2012).

16 Gao, J. et al. Integrative analysis of complex cancer genomics and clinical profiles using the cBioPortal. Sci Signal 6, pl1, doi:10.1126/scisignal.2004088 (2013).

17 Johnson, D. E. et al. Head and neck squamous cell carcinoma. Nat Rev Dis Primers 6, 92, doi:10.1038/s41572-020-00224-3 (2020).

18 Zhuang, L. et al. Depletion of Nsd2-mediated histone H3K36 methylation impairs adipose tissue development and function. Nat Commun 9, 1796, doi:10.1038/s41467-018-04127-6 (2018).

19 Farhangdoost, N. et al. Chromatin dysregulation associated with NSD1 mutation in head and neck squamous cell carcinoma. Cell Rep 34, 108769, doi:10.1016/j.celrep.2021.108769 (2021).

20 Rajagopalan, K. N. et al. Depletion of H3K36me2 recapitulates epigenomic and phenotypic changes induced by the H3.3K36M oncohistone mutation. Proc Natl Acad Sci U S A 118, doi:10.1073/pnas.2021795118 (2021).

21 Bai, Y. et al. The Balance between Differentiation and Terminal Differentiation Maintains Oral Epithelial Homeostasis. Cancers (Basel) 13, doi:10.3390/cancers13205123 (2021).

22 Muraoka-Cook, R. S. et al. The intracellular domain of ErbB4 induces differentiation of mammary epithelial cells. Mol Biol Cell 17, 4118–4129, doi:10.1091/mbc.e06-02-0101 (2006).

23 Ochiai, H. et al. BMP4 and FGF strongly induce differentiation of mouse ES cells into oral ectoderm. Stem Cell Res 15, 290–298, doi:10.1016/j.scr.2015.06.011 (2015).

24 Gong, S. G., Mai, S., Chung, K. & Wei, K. Flrt2 and Flrt3 have overlapping and non-overlapping expression during craniofacial development. Gene Expr Patterns 9, 497–502, doi:10.1016/j.gep.2009.07.009 (2009).

25 Bian, Y. et al. Loss of TGF-beta signaling and PTEN promotes head and neck squamous cell carcinoma through cellular senescence evasion and cancer-related inflammation. Oncogene 31, 3322–3332, doi:10.1038/onc.2011.494 (2012).

26 Morad, G., Helmink, B. A., Sharma, P. & Wargo, J. A. Hallmarks of response, resistance, and toxicity to immune checkpoint blockade. Cell 184, 5309–5337, doi:10.1016/j.cell.2021.09.020 (2021).

27 Fiore, P. F. et al. Interleukin-15 and cancer: some solved and many unsolved questions. J Immunother Cancer 8, doi:10.1136/jitc-2020-001428 (2020).

28 Musselmann, K. et al. Salivary gland gene expression atlas identifies a new regulator of branching morphogenesis. J Dent Res 90, 1078–1084, doi:10.1177/0022034511413131 (2011).

29 Veniaminova, N. A. et al. Niche-Specific Factors Dynamically Regulate Sebaceous Gland Stem Cells in the Skin. Dev Cell 51, 326–340 e324, doi:10.1016/j.devcel.2019.08.015 (2019).

30 Tecles, F. et al. Total esterase activity in human saliva: Validation of an automated assay, characterization and behaviour after physical stress. Scand J Clin Lab Invest 76, 324–330, doi:10.3109/00365513.2016.1163417 (2016).

31 Saitou, M. et al. Functional Specialization of Human Salivary Glands and Origins of Proteins Intrinsic to Human Saliva. Cell Rep 33, 108402, doi:10.1016/j.celrep.2020.108402 (2020).

32 Delporte, C. & Steinfeld, S. Distribution and roles of aquaporins in salivary glands. Biochim Biophys Acta 1758, 1061–1070, doi:10.1016/j.bbamem.2006.01.022 (2006).

33 Motegi, K., Azuma, M., Tamatani, T., Ashida, Y. & Sato, M. Expression of aquaporin-5 in and fluid secretion from immortalized human salivary gland ductal cells by treatment with 5-aza-2’-deoxycytidine: a possibility for improvement of xerostomia in patients with Sjogren’s syndrome. Lab Invest 85, 342–353, doi:10.1038/labinvest.3700234 (2005).

34 Zhou, H. D. et al. Tissue distribution of the secretory protein, SPLUNC1, in the human fetus. Histochem Cell Biol 125, 315–324, doi:10.1007/s00418-005-0070-4 (2006).

35 Woods, R. S. R. et al. Cytokeratin 7 and 19 expression in oropharyngeal and oral squamous cell carcinoma. Eur Arch Otorhinolaryngol 279, 1435–1443, doi:10.1007/s00405-021-06894-3 (2022).

36 An, Q. et al. KRT7 promotes epithelialmesenchymal transition in ovarian cancer via the TGFbeta/Smad2/3 signaling pathway. Oncol Rep 45, 481–492, doi:10.3892/or.2020.7886 (2021).

37 Yan, G. et al. Single-cell transcriptomic analysis reveals the critical molecular pattern of UV-induced cutaneous squamous cell carcinoma. Cell Death Dis 13, 23, doi:10.1038/s41419-021-04477-y (2021).

38 Zulijani, A., Dekanic, A., Cabov, T. & Jakovac, H. Metallothioneins and Megalin Expression Profiling in Premalignant and Malignant Oral Squamous Epithelial Lesions. Cancers (Basel) 13, doi:10.3390/cancers13184530 (2021).

39 Irie, Y., Mori, F., Keung, W. M., Mizushima, Y. & Wakabayashi, K. Expression of neuronal growth inhibitory factor (metallothionein-III) in the salivary gland. Physiol Res 53, 719–723 (2004).

40 Fang, D. et al. The histone H3.3K36M mutation reprograms the epigenome of chondroblastomas Science 352, 1344–1348, doi:10.1126/science.aae0065 (2016).

41 Streubel, G. et al. The H3K36me2 Methyltransferase Nsd1 Demarcates PRC2-Mediated H3K27me2 and H3K27me3 Domains in Embryonic Stem Cells. Mol Cell 70, 371–379 e375, doi:10.1016/j.molcel.2018.02.027 (2018).

42 Ezhkova, E. et al. Ezh2 orchestrates gene expression for the stepwise differentiation of tissue-specific stem cells. Cell 136, 1122–1135, doi:10.1016/j.cell.2008.12.043 (2009).

43 Brumbaugh, J. et al. Inducible histone K-to-M mutations are dynamic tools to probe the physiological role of site-specific histone methylation in vitro and in vivo. Nat Cell Biol 21, 1449–1461, doi:10.1038/s41556-019-0403-5 (2019).

44 Meisel, C. T., Pagella, P., Porcheri, C. & Mitsiadis, T. A. Three-Dimensional Imaging and Gene Expression Analysis Upon Enzymatic Isolation of the Tongue Epithelium. Front Physiol 11, 825, doi:10.3389/fphys.2020.00825 (2020).

45 Egolf, S. et al. LSD1 Inhibition Promotes Epithelial Differentiation through Derepression of Fate-Determining Transcription Factors. Cell Rep 28, 1981–1992 e1987, doi:10.1016/j.celrep.2019.07.058 (2019).

46 Dobin, A. et al. STAR: ultrafast universal RNA-seq aligner. Bioinformatics 29, 15–21, doi:10.1093/bioinformatics/bts635 (2013).

47 Love, M. I., Huber, W. & Anders, S. Moderated estimation of fold change and dispersion for RNA-seq data with DESeq2. Genome Biol 15, 550, doi:10.1186/s13059-014-0550-8 (2014).

48 Yang, Z., Guan, B., Men, T., Fujimoto, J. & Xu, X. Comparable molecular alterations in 4-nitroquinoline 1-oxide-induced oral and esophageal cancer in mice and in human esophageal cancer, associated with poor prognosis of patients. In Vivo 27, 473–484 (2013).

49 Egolf, S. et al. MLL4 mediates differentiation and tumor suppression through ferroptosis. Sci Adv 7, eabj9141, doi:10.1126/sciadv.abj9141 (2021).

50 Aziz, S. et al. Evaluation of Tumor Cell Proliferation by Ki-67 Expression and Mitotic Count in Lymph Node Metastases from Breast Cancer. PLoS One 11, e0150979, doi:10.1371/journal.pone.0150979 (2016).

