## Extended Data Table 2 and 3 for "H3K36M provokes cellular plasticity to drive aberrant glandular formation and squamous carcinogenesis"

**Data on 4NQO induced oral and esophageal cancer development in male mice**

| Treatment groups (Male) | | Initial weight (g) | Final weight (g) | Site (Number) | N of animal affected | H* | D* | T* |
|---|---|---|---|---|---|---|---|---|
| WT_Water | 5 | 25.22 ± 1.69 | 34.0 ± 1.40 | Tongue (5) | 0 | | | |
| | | | | Esophagus (5) | 0 | | | |
| K36M_Water | 5 | 21.04± 2.69 | 30.7 ± 2.95 | Tongue (5) | 2 | 2 | | |
| | | | | Esophagus (4) | 2 | 2 | | |
| WT_4NQO | 5 | 24.41 ± 0.94 | 30.5 ± 1.26 | Tongue (5) | 5 | 4 | 1 | |
| | | | | Esophagus (4) | 4 | 3 | 1 | |
| K36M_4NQO | 6 | 22.15 ± 1.92 | 22.45 ± 3.70 | Tongue (6) | 6 | 2 | 3 | 1 |
| | | | | Esophagus (5) | 5 | 2 | 2 | 1 |

Extended Data Table 3

**Data on 4NQO induced oral and esophageal cancer development in female mice**

| **Treatment groups (Female)** | **N of mice** | **Initial weight (g)** | **Final weight (g)** | **Site (Number)** | **N of animal affected** | **H*** | **D*** | **T*** |
|---|---|---|---|---|---|---|---|---|
| WT_Water | 8 | 18.50 ± 1.15 | 25.95 ± 1.09 | Tongue (8) | 0 | | | |
| | | | | Esophagus (7) | 0 | | | |
| K36M_Water | 7 | 17.37± 2.72 | 23.92 ± 3.05 | Tongue (7) | 5 | 5 | | |
| | | | | Esophagus (7) | 5 | 5 | | |
| WT_4NQO | 7 | 18.35 ± 1.33 | 22.16 ± 2.31 | Tongue (7) | 6 | 5 | 1 | |
| | | | | Esophagus (6) | 5 | 3 | 2 | |
| K36M_4NQO | 9 | 16.9 ± 1.92 | 17.81 ± 3.08 | Tongue (9) | 9 | 2 | 3 | 4 |
| | | | | Esophagus (9) | 8 | 2 | 6 | |
